# Long non-coding RNA G23Rik attenuates fasting-induced lipid accumulation in the liver

**DOI:** 10.1101/2022.04.11.487900

**Authors:** Donghwan Kim, Bora Kim, Chad N. Brocker, Kritika Karri, David J. Waxman, Frank J. Gonzalez

**Affiliations:** Laboratory of Metabolism, Center for Cancer Research, National Cancer Institute, National Institutes of Health, Bethesda, Maryland 20892; Department of Biology and Bioinformatics Program, Boston University, Boston, Massachusetts 02215

**Keywords:** PPARA, LncRNA, G23Rik, fasting, lipid accumulation

## Abstract

Peroxisome proliferator-activated receptor α (PPARα) is a key mediator of lipid metabolism and metabolic stress in the liver. A recent study revealed that PPARα-dependent long non-coding RNAs (lncRNAs) play an important role in modulating metabolic stress and inflammation in the livers of fasted mice. Here hepatic lncRNA *3930402G23Rik* (*G23Rik*) was found to have active peroxisome proliferator response elements (PPREs) within its promoter and is directly regulated by PPARα. Although *G23Rik* RNA was expressed to varying degrees in several tissues, the PPARα-dependent regulation of this lncRNA was only observed in the liver. Pharmacological activation of PPARα induced PPARα recruitment at the *G23Rik* promoter and a pronounced increase in hepatic *G23Rik* lncRNA expression. A *G23Rik*-null mouse line was developed to further characterize the function of this lncRNA in the liver. *G23Rik*-null mice were more susceptible to hepatic lipid accumulation in response to acute fasting. Histological analysis further revealed a pronounced buildup of lipid droplets and a significant increase in neutral triglycerides and lipids as indicated by enhance oil red O staining of liver sections. Hepatic cholesterol, non-esterified fatty acid, and triglyceride levels were significantly elevated in *G23Rik*-null mice and associated with induction of the lipid-metabolism related gene *Cd36*. These findings provide evidence for a lncRNA dependent mechanism by which PPARα attenuates hepatic lipid accumulation in response to metabolic stress through lncRNA *G23Rik* induction.

**Key points:**

1. 3930402G23Rik (G23Rik) is a liver-enriched long non-RNA.
2. G23Rik is a PPARA target gene.
3. Loss of G23Rik increases hepatic lipid accumulation during fasting

## Introduction

Peroxisome proliferator-activated receptor α (PPARα) is a ligand activated nuclear receptor that is a key mediator of lipid metabolism in liver. PPARα is a therapeutic target for metabolic diseases associated with hyperlipidemia, with the potential for treatment of type 2 diabetes, and obesity 1. Notably, the fibrate class of hyperlipidemic drugs includes PPARα agonists that also act as potent anti-inflammatory agents in humans ^2, 3^. In contrast, chronic activation of PPARα by the potent agonist WY14643 promotes hepatocyte proliferation and hepatocellular carcinogenesis in mice 4. The role of PPARα is extensive and diverse and the regulatory mechanisms underlying how PPARα suppresses inflammation are still not well defined.

Long non-coding RNAs (lncRNAs) are non-coding RNAs with lengths exceeding 200 nucleotides that regulate cellular functions through multiple mechanism 5. LncRNAs play important roles in developmental regulation and cell signaling, and are many are highly tissue and cell state-specific; moreover, the number of expressed non-coding RNAs often increases with the complexity and differentiation status of the cell 6. Thus, the lncRNA HISLA, expressed in tumor-associated macrophages, promotes glycolysis and decreases apoptosis by inhibiting hypoxia-inducible factor (HIF)1α in tumor cells 7, while lncRNA-FEZF1-AS1 and lncRNA-00184 directly regulate glucose metabolism by inducing target genes associated with glucose metabolism ^8, 9^. LncRNAs also play a regulatory role in lipid metabolism in macrophages associated with inflammation-related diseases 10. Notably, the liver-enriched PPARα regulated lncRNA *Gm15441* was found to modulate metabolic stress by inhibiting the *Txnip*-mediated *NLRP3* inflammasome pathway in liver 11. Other PPARα inducible lncRNAs may therefore also play important roles in metabolic remodeling and anti-inflammatory actions in the liver and possibly other tissues.

*3930402G23Rik (G23Rik) (lnc_inter_c8_6744* ^11, 12^) was identified as a highly upregulated lncRNA in livers of mice treated with the PPARA agonist WY14643. The physiological function of this lncRNA in the liver is unknown. Gene expression studies identified *G23Rik* as differentially regulated in several tissues. *G23Rik* is enriched in brown fat when compared to white fat and expressed at higher levels in differentiated brown adipocytes than 3T3-L1 adipocytes 13. This expression pattern paralleled the expression of brown fat lncRNA 1 (*Blnc1*). *Blnc1* was shown to promote brown and beige adipocyte differentiation and function 14, suggesting *G23rik* may play a role in this process. *G23Rik* was also identified as one of the top 50 differentially regulated genes in the pancreatic islet cells of mice lacking the short chain fatty acid receptor free fatty acid receptor-3 (*Ffar3*) which regulates glucose stimulated insulin secretion 15. A related short chain fatty acid receptor, *Ffar4*, is downregulated ten-fold in liver after PPARA activation (GSE132385 and GSE132386) 11. Together, these expression data support *G23Rik*’s potential role in metabolic processes.

In this study, the mechanisms by which the liver-enriched and PPARα-inducible lncRNA *G23Rik* influences lipid accumulation during the physiological response to fasting was investigate. ChIP-seq datasets were analyzed for PPARα binding sites within the *G23Rik* promoter, and *G23Rik* regulatory elements were confirmed using luciferase reporter gene assays and PPARα ChIP-PCR analysis. A *G23Rik* knockout mouse model was developed by employing CRIPSR/Cas9-mediated gene editing and the subsequent deletion of exon 3 (*G23Rik*^-/-^). *G23Rik*^-/-^ mice were treated with a PPARα agonist or fasted to determine how loss of *G23Rik* impacts hepatic damage in response to both pharmacological stress (i.e., WY14643-treatment) and physiological metabolic stress (i.e., fasting). Hepatic lipid droplet and lipid accumulation were analyzed in fasted *G23Rik*^-/-^ mice and compared to fasted *G23Rik*+/+ mice. Lipid accumulation-related gene expression was also significantly altered in livers of fasted *G23Rik*^-/-^ mice. Taken together, these findings support a regulatory mechanism involving PPARα targeting lncRNA *G23Rik* during fasting.

## Material and methods

### Mouse models

Male 8- to 12-week-old mice were used for all studies. All mouse lines were on the C57BL/6J background and maintained on a grain-based control diet (NIH-31). Mice were housed in light and temperature-controlled rooms and were provided with water and pelleted chow *ad libitum*. For pharmacological studies, mice were fed NIH-31 diet or a matched NIH-31 diet containing 0.1% WY14643 for 48 hrs. When monitoring the time dependence of gene expression responses, WY14643 was dissolved in 1% carboxymethyl cellulose (CMC) solution and orally administered (50 mg/kg in 200 μl) by gavage. At the indicated time points, mice were killed by CO2 asphyxiation and tissues harvested, quick frozen in liquid N2 and stored at -80 oC until processing. For physiological studies, food was removed for 24 h starting shortly after the onset of the light cycle, and endpoints were collected at the same time the following day. Mice were euthanized and tissue samples were harvested for further analysis. Blood was collected by venipuncture of the caudal vena cava. All animal experiments were performed in accordance with the Association for Assessment and Accreditation of Laboratory Animal Care international guidelines and approved by the National Cancer Institute Animal Care and Use Committee.

### Generation of G23Rik-null mice

SAGE Laboratories (Cambridge, UK) provided design and construction services for the CRISPR/Cas9 gene targeting technologies used to create a *G23Rik*-null (*G23Rik*^-/-^) mouse line. Cleavage at the insertion sites excises exon 3 of *G23Rik* and abolishes *G23Rik* expression, resulting in a conventional whole-body knockout mouse line. Microinjection-ready sgRNA and *Cas9* mRNA were purchased from SAGE Laboratories. sgRNA and Cas9 mRNA were then injected into C57BL/6J mouse embryos by the Transgenic Mouse Model Laboratory at the National Cancer Institute (Fredrick, MD) using the manufacturer’s recommended protocol. Founder animals were genotyped using primer sets listed in **Table S1**, and all modifications were confirmed by targeted sequencing. Homozygous mice were then backcrossed ten times into the C57BL/6J background.

### Mouse genotyping

Genomic DNA was extracted from tail snips using the Sigma REDExtract-N-Amp Kit. Genotyping was performed following the manufacturer’s protocol with the following modifications. Twenty-five µl of E buffer and 7 ul of TPS buffer were added and stored room temperature at 10 min. Then tails were heated at 95 oC for 5 min and 25 µl of N buffer was added. Polymerase chain reaction was used to determine the genotype using primers shown in **Table S1**. Amplicons of lengths 900 and 1200 bp were detected in wild-type and *G23Rik*-null mice, respectively.

### RNA isolation, library construction, and RNA sequencing

RNeasy Plus Mini Kits (Qiagen, Valencia, CA, USA) were used to extract total RNA from livers from four different treatment and control groups, namely wild-type (*G23Rik*^+/+^) and *G23Rik*^−/−^ mice fed a control diet a matching diet containing WY14643 for 48 h. All mice were killed between 1 and 3 PM. RNAs were extracted from n = 9 to 15 independent livers per group and RNA quality assessed using a TapeStation 4200 instrument (Agilent, Santa Clara, CA, USA). High quality RNA samples (RIN > 9.0) were pooled, as described below, and used to construct stranded RNA-seq libraries from polyA-selected total liver RNA using an Illumina stranded TruSeq mRNA Prep Kit (Illumina, San Diego, CA, USA). Three pooled RNA samples were prepared for each of the four treatment groups, with each pool comprised of n = 3-5 individual liver samples. The libraries were subjected to paired-end sequencing using an Illumina HiSeq 2500 instrument (Illumina) at the NCI-CCR sequencing facility (Frederic, USA) at a depth of 30-42 million read pairs for each of the 12 RNA-seq libraries. RNA-seq data was analyzed using an in-house custom RNA-seq pipeline 16. Briefly, sequence reads were aligned to mouse genome build mm9 (NCBI 37) using TopHat (version 2.2.5.0) 17. FeatureCounts 18 was used to count sequence reads mapping to the union of the exonic regions in all isoforms of a given gene (collapsed exon counting), and differential expression analysis was conducted using the Bioconductor package EdgeR 19 to identify differentially expressed genes for each indicated comparison, using the following cutoffs: p-value < 0.05 an |fold-change| > 2.0. Raw RNA-seq files and processed data files are available at GEO (https://www.ncbi.nlm.nih.gov/geo/), accession # GSE197885.

### Cell culture and transfection

Primary hepatocytes were isolated from C57BL/6J mice as previously reported 20 and seeded at a density of 2 × 105 cells on collagen-coated 12-well plates (Becton Dickinson and Company, Franklin Lakes, NJ) in Williams’ Medium E (Thermo-Fisher Scientific, Waltham, MA) supplemented with 5% FBS and penicillin/streptomycin/amphotericin B solution (Gemini Bio-products, West Sacramento, CA).

### Histological staining

Fresh liver tissue was immediately fixed in 10% phosphate-buffered formalin for 24 h and then processed in paraffin blocks. Four-micrometer sections were used for H&E staining. For Oil Red O (ORO) staining, fresh liver tissue was placed into a cryomold and filled with OCT Compound (Tissue-Tek), then transferred to a beaker of isopentane prechilled in liquid nitrogen. Sections were processed by HistoServ, Inc. (Germantown, MD). Unstained slides were fixed in 10% neutral buffered formalin (Sigma), rinsed in water and then transferred to 100% propylene glycol (Sigma). Slides were then stained in 0.5% ORO solution (Sigma), washed in 85% propylene glycol, then water. The slides were then counterstained with Carazzi’s hematoxylin and rinsed in water. Glycerin jelly was used to mount slides. Slide imaging was performed using a Keyence BZ-X700 series all-in-one microscope with 20× objectives, 200× magnification.

### Serum biochemistry

Blood was collected from mice and transferred to BD Microtainer Serum Separator Tubes (Becton Dickinson, Franklin Lakes, NJ). Serum was flash frozen in liquid nitrogen and stored at -80C. Serum chemistry analysis for total cholesterol (CHOL), non-esterified fatty acids (NEFA), and triglycerides (TG) was performed using Wako Clinical Diagnostics kits (WakoUSA, Richmond, VA). Serum alanine aminotransferase (ALT) and aspartate aminotransferase (AST) levels were measured using Catachem VETSPEC Kits as recommended by the manufacturer (Catachem, Oxford, CT).

### Quantitative reverse transcription PCR assays

Total RNA was isolated from fresh mouse liver, mouse primary hepatocytes, and Hepa-1 cells using TRIzol Reagent (Thermo-Fisher Scientific, Waltham, MA, USA) and quantified using a NanoDrop Spectrophotometer (NanoDrop Products, Wilmington, DE, USA). Total RNA (1 μg) was reverse transcribed using qScript cDNA Synthesis Kit (VWR, Houston, TX, USA). qRT-PCR analysis was performed using SYBR Green qPCR Master Mix (VWR). Primers were designed for gene specificity and to cross exon-exon junctions using Primer-BLAST (www.ncbi.nlm.nih.gov/tools/primer-blast/) and purchased from IDT DNA Technologies (Coralville, IA, USA) (**Table S1**). qRT-PCR experiments were designed and performed according to Minimum Information for Publication of Quantitative Real-Time PCR Experiments (MIQE) guidelines. Results are normalized to actin expression. Values given are fold over control or relative expression value, as appropriate, and were calculated using the 2ΔCt QPCR calculation method.

### Luciferase reporter assays

For luciferase assays, pSG5-PPARα (mouse) and pSG5-RXRα (mouse) were used for transcription factor expression. Custom GeneBlocks (IDT DNA) were synthesized containing the predicted PPRE sites for *G23Rik* (**Table S1**). GeneBlocks were digested and purified using a Qiagen PCR Purification Kit (Qiagen, Valencia, CA) and cloned into the pGL4.11 for PPRE reporter constructs (Promega, Madison, WI) using a BioRad Quick Ligation Kit (BioRad, Hercules, CA, USA). Prior to performing assays, all constructs were confirmed by sequencing at the NCI Center for Cancer Research Genomics Core. The phRL-TK renilla luciferase construct was used as a control to normalize for transfection efficiency. Primary hepatocytes were seeded into 12-well plates (4 × 104 cells/well). PPRE reporter constructs were co-transfected into hepatocytes with PPARα and RXRα expression vectors. PPRE-luc plasmid containing an *Acox1* PPRE site repeat was used as a positive control 21. Empty pGL4.11 plasmids were used as negative controls. Plasmids were transfected using Lipofectamine 3000 Reagent (Thermo-Fisher Scientific). Luciferase activities were measured and plotted relative to lysate protein concentrations using the Promega Dual Luciferase Reporter (Promega) assays according to the manufacturer’s protocol. Measurements were taken on a Veritas microplate luminometer (Turner Biosystems, Sunnyvale, CA, USA).

### Chromatin immunoprecipitation

Chromatin was prepared from hepatocytes for ChIP assays as previously described. Cells were fixed with 4% paraformaldehyde for 15 min, then glycine was added to a final concentration of 0.125 M and incubated for 10 min before harvesting. Chromatin was sonicated using a Bioruptor Pico instrument (Diagenode, Denville, NJ, USA). Chromatin preparations were subjected to ChIP using a ChIP-IT High Sensitivity Kit and Protein G Agarose Prepacked Columns (Active Motif, Carlsbad, CA, USA) using a PPARα (Abcam Ab24509) antibody. Normal rabbit IgG (Cell Signaling Technologies #2729S) and Histone H3 antibody (Cell Signaling Technologies #4620) were used as negative and positive controls, respectively. DNA was purified and concentrated using MinElute Reaction Cleanup columns (Qiagen). qRT-PCR and conventional PCR were performed using 2 μl of ChIP DNA samples from the 50 μl of purified samples using gene-specific primers (**Table S1**). Cycle threshold (Ct) values of ChIP and input samples were calculated and presented as fold change values.

### Sample preparation and analysis for lipid profiling

Liver samples (∼15 mg) were weighed and homogenized in 56 uL/(mg tissue weight) of water-acetonitrile (3:4 v/v) in a Precellys 24 bead homogenizer (Bertin, Rockville, MD). Chloroform was added with 16 μl/(mg tissue weight), and the mixture was vortexed for 1 min. Samples were centrifuged at 3,000g for 20 min at 4 °C to separate the phases. Organic phases were transferred to glass tubes to which chloroform containing butylated hydroxytoluene at a final concentration of 1 mM. The sample was vortexed, then evaporated under nitrogen, and stored at 80 °C until just before analysis. The dried extracts were dissolved in 20 uL/(mg tissue weight) of isopropanol-methanol-chloroform (4:2:1 v/v/v) and transferred to ultraperformance liquid chromatography (UPLC) injection vials. Samples were analyzed by UPLC-electrospray ionization-quadrupole time-of-flight mass spectrometry (UPLC-ESI-QTOFMS) using an Acquity UPLC CSH C18 1.7 μm (2.1 × 100 mm) under the following conditions: solvent A, 10 mM ammonium formate. 0.1% formic acid acetonitrile/water (60/40); solvent B, 10 mM ammonium formate, 0.1% formic acid in IPA/acetonitrile (90/10); 60% solvent A for initial, to 57% solvent A at 2 min, to 50% solvent A at 2.1 min, to 46% solvent A at 12 min, to 30% solvent A at 12.1 min, to 1% solvent A at 18 min, to 60% solvent A at 18.1 min, then equilibriated at initial conditions for 1.9 min. Flow rate was 0.4 ml/min, and column temperature was maintained at 55 °C. Mass spectrometry was conducted in positive (ESI+) and negative (ESI–) modes using a Synapt G2S Q-TOF MS system (Waters, Milford, MA), scanning 100–1200 amu. Capillary voltage was 2.82 kV, source temperature was 150 °C, sampling cone was 41V, and desolvation gas flow was 950 l/h at 500 °C. Pooled sample was injected every 4 samples to monitor instrument stability.

### Data and statistical analyses

PPARα ChIP-seq data was downloaded from the NCBI Gene Expression Omnibus (GSE61817). CHIP-seq data was uploaded to the Galaxy public server at usegalaxy.org to analyze the data 22. Data were converted to bigwig file format using Galaxy tools and then visualized using Integrated Genome Browser (version 9.0.0). All results are expressed as means ± SD. Significance was determined by t-test or one-way ANOVA with Bonferroni correction using Prism 7.0 software (GraphPad Software, La Jolla, CA, USA). The lipidomics raw data were imported, deconvoluted, normalized and reviewed using Progenesis QI software (Waters). A multivariate data matrix was generated through centroiding, deisotoping, filtering, and integration. The data matrix was further analyzed using SIMCA version 16.0.2 (Umetrics, Kinnelon, NJ). Principle component analysis (PCA) was used to examine the separation between group and confirm the stability and reproducibility of the data. A P value less than 0.05 was considered significant as indicated in the figure legends.

## Results

### LncRNA G23Rik is PPAR *α-*dependent and induction is liver-specific

Gene expression profiling by RNA-seq analysis identified many lncRNAs whose expression responds to activation of PPARα by WY14643 in mouse liver (GSE132386), including *Gm15441*, a highly liver specific PPARα target gene that modulates liver stress induced by WY14643 and fasting 11. Expression of another transcript, *G23Rik*, was significantly elevated in WY14643-treated *Ppara*^+/+^ liver but was completely abolished in WY14643-treated *Ppara*^-/-^ liver 11. Further studies were performed to characterize the tissue specificity and PPARα dependence of *G23Rik* expression.

Basal expression levels of *G23Rik* were assayed across six mouse tissues. In the absence of WY14643 treatment, *G23Rik* was most highly expressed in muscle and brown adipose tissue (BAT) (**Fig. 1A**). To assess the impact of PPARα activation on *G23Rik* expression and its tissue specificity, wild-type mice (*Ppara*^+/+^) and *Ppara*-null mice (*Ppara*^-/-^) were treated with WY14643 for 48 h, and seven tissues then harvested for analysis. *G23Rik* mRNA levels were strongly increased by WY14643 in a liver-specific manner in *Ppara*^+/+^ mice; in contrast, no induction of *G23rik* mRNA was seen in livers of WY14643-treated *Ppara*^-/-^ mice (**Fig. 1B**). No significant induction was seen in the six other tissues, consistent with the tissue specificity of Ppara expression and activity (**Fig. 1B**). *G23Rik* induction in liver was rapid, with maximum expression seen 6 h after a single gavage dose of WY14643 (**Fig. 1C**). Thus, *G23Rik* is highly and specifically inducible by PPARα in liver.

**Fig. 1.**
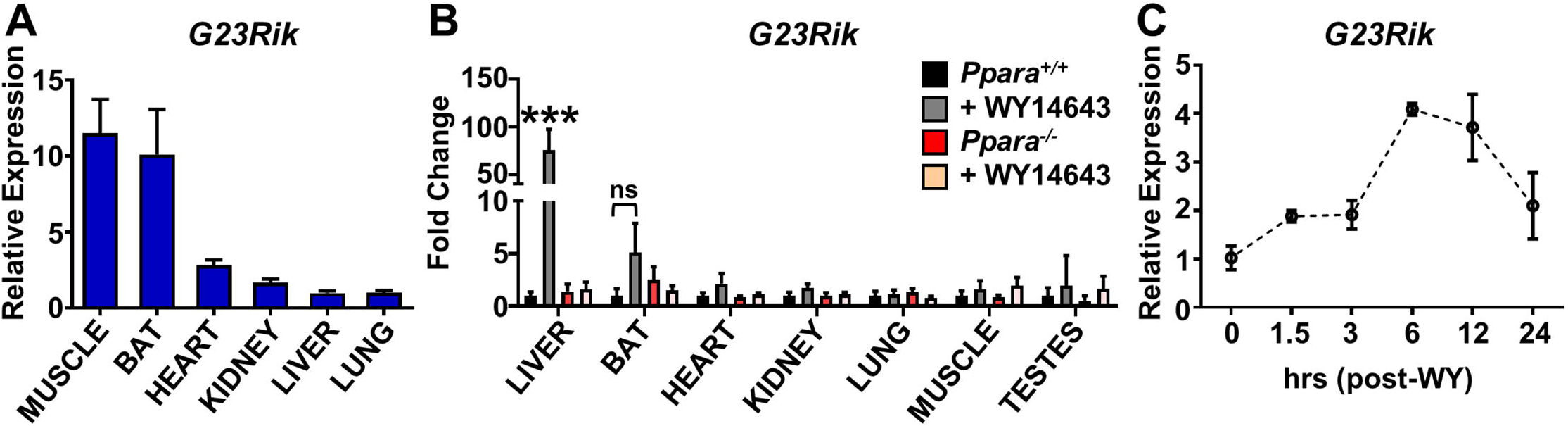
LncRNA *G23Rik* is PPARA dependent. **(A)** Relative basal *G23Rik* expression in select tissues. **(B)** *G23Rik* expression in tissues from *Ppara*^+/+^ and *Ppara*^-/-^ mice treated with WY-14643 (via diet) for 48 h. **(C)** Time course for changes in expression of *G23Rik* over a 24 h period following treatment with WY14643 (50 mg/kg in 200 μl by gavage). The maximum response of *G23Rik* was seen at 6 h. **P < 0.01; ***P < 0.001

### PPARα directly regulates lncRNA G23Rik

PPARα ChIP-seq data (GSE61817) revealed one major ChIP-seq peak indicating PPARα binding at a site upstream of *G23Rik* (**Suppl. Fig. S1**). Analysis of genomic sequences upstream of *G23Rik* using Genomatix MatInspector (Genomatix, Munich, Germany) identified five upstream regions, designated A to E, that contain six peroxisome proliferator response elements (PPREs), all within 5 kilobase (kb) of the *G23Rik* transcriptional start site (**Fig. 2A**). These five *G23Rik* upstream sequences were synthesized and cloned into the pGL4.11 reporter, and luciferase reporter gene assays were performed to determine the functionality of PPARα binding to these *G23Rik* promoter sequences. A PPRE-luciferase construct containing an *Acox1* PPRE repeat was used as a positive control, and an empty pGL4.11 plasmid was used as a negative control. Luciferase activity was significantly elevated in primary mouse hepatocytes transfected with upstream region C, but not the other four pGL4.11 constructs, consistent with direct regulation by PPARα (**Fig. 2B**). PPARα binding was also assessed by ChIP assays using a polyclonal PPARα antibody and chromatin isolated from livers of wild-type and *Ppara*^-/-^ mice fed either chow diet or a diet-containing WY-14643. Enrichment of PPARα binding to PPREs of the known PPARα target genes *Acot1* encoding acyl-CoA thioesterase 1 and *Acox1* encoding acyl-Co-A oxidase 1 was determined by comparing binding to liver chromatin from wild-type mice fed chow diet vs. mice fed WY14643-containing diet (**Fig. 2C**). Livers from *Ppara*^-/-^ mice were used as a negative control to identify non-specific binding. PPARα binding was significantly enriched at the genomic region amplified using primers covering *G23Rik* region C, but not at the other four *G23Rik* upstream regions (**Fig. 2C**), which were transcriptionally inactive (**Fig. 2B**). Binding at site C was significantly increased by WY14643 treatment, indicating increased PPARα recruitment to these sites. No PPARα binding was seen with chromatin from *Ppara*- /- mice on the control diet (**Fig. 2C**). We conclude that PPARα regulates *G23Rik* transcription by direct binding of PPARα to a PPRE site 2.1 kb upstream of the *G23Rik* transcriptional start site.

**Fig. 2.**
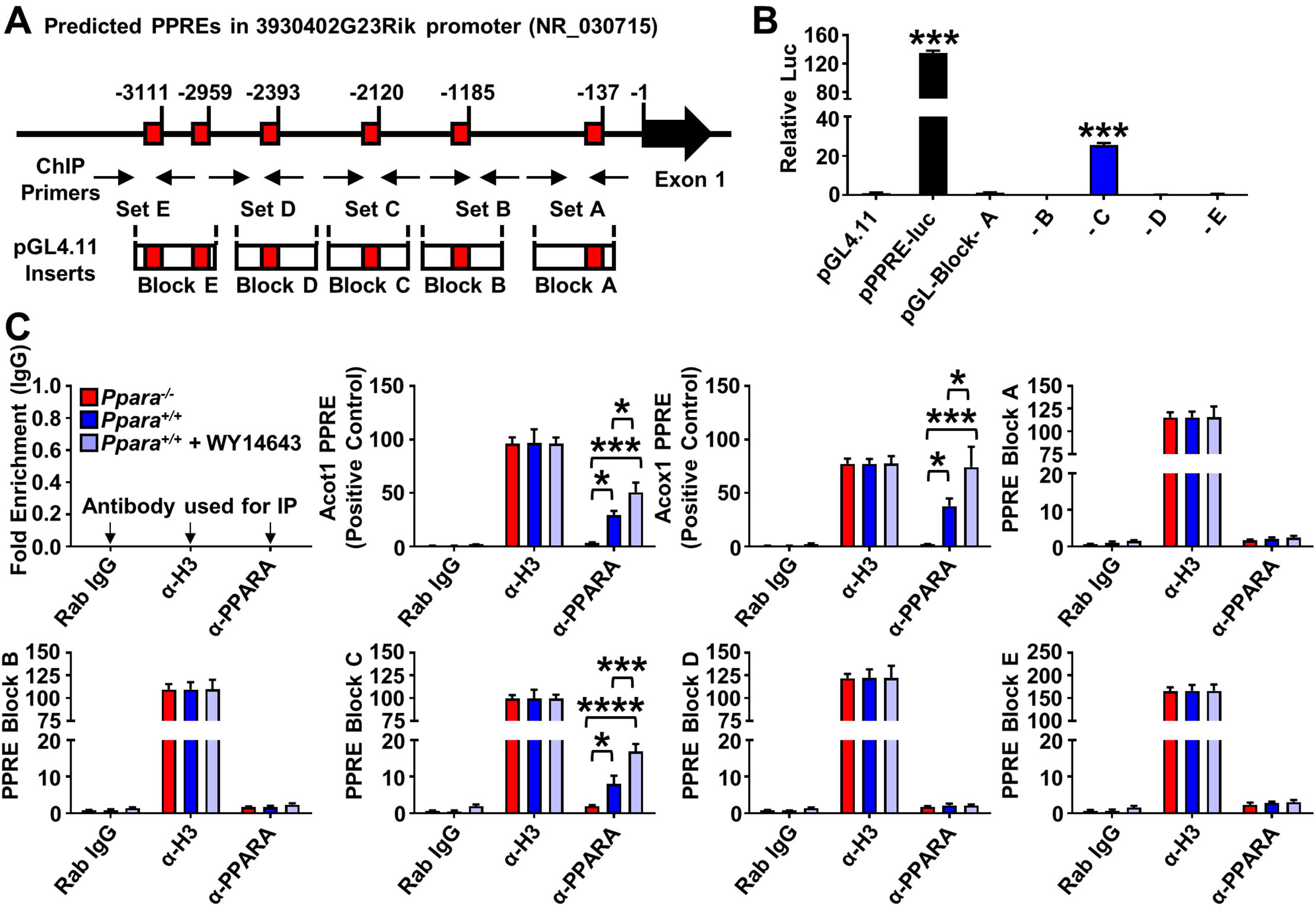
LncRNA *G23Rik* is a direct PPAR α target gene. **(A)** Schematic representation of seven PPRE sequences found within the *G23Rik* promoter (−5 kb). ChIP primer binding sites and reporter gene construct inserts are shown. **(B)** Luciferase-based reporter assays identified five functional PPREs within the *G23Rik* promoter, based on n = 3 replicates. **(C)** PPARα ChIP assays of PPRE binding in liver samples from *Ppara*^+/+^ and *Ppara*^-/-^ mice treated with WY14643. Experiments were performed with at least four different livers. Rabbit IgG and antibody to histone protein H3 were used as negative and positive controls, respectively. Each data point represents the mean ± SD for n = 5 liver samples. *P < 0.05; ***P < 0.001; ****P < 0.0001.

### Generation of a G23Rik knockout mouse

Next, a conventional whole-body *G23Rik* knockout mouse model was generated using CRISPR/Cas9 to introduce cleavage sites flanking *G23Rik* exon 3 (*G23Rik*^-/-^, **Fig. 3A**). All mouse lines were then bred to give a pure C57BL/6J background. The genotyping scheme generates an approximately 900 bp band in *G23Rik* wild-type mice (*G23Rik*^+/+^) and a 1200 bp band in *G23Rik* knockout mice (*G23Rik*^-/-^) (**Fig. 3B**). WY14643 treatment induced *G23Rik* in *G23Rik*^+/+^ and *G23Rik*+/- mice, but not in *G23Rik*^-/-^ mice (**Fig. 3C**), validating *G23Rik*^-/-^ mice as an effective knockout mouse model. Pronounced hepatomegaly was found after WY14643 treatment in both *G23Rik*^+/+^ and *G23Rik*^-/-^ mice, as indicated by the pronounced swelling of hepatocytes seen by H&E staining (**Fig. 3D**), and by the significant increase in liver size (**Fig. 3E**, liver index). Further, there was no discernable difference in liver index and total body weight loss between *G23Rik*^+/+^ and *G23Rik*^-/-^ mice on a WY14643 diet (**Fig. 3E**). Triglyceride (TG) and cholesterol levels in liver were decreased by WY14643 feeding, but with no difference in *G23Rik*^+/+^ compared to *G23Rik*^-/-^ mice (**Fig. 3E**).

**Fig. 3.**
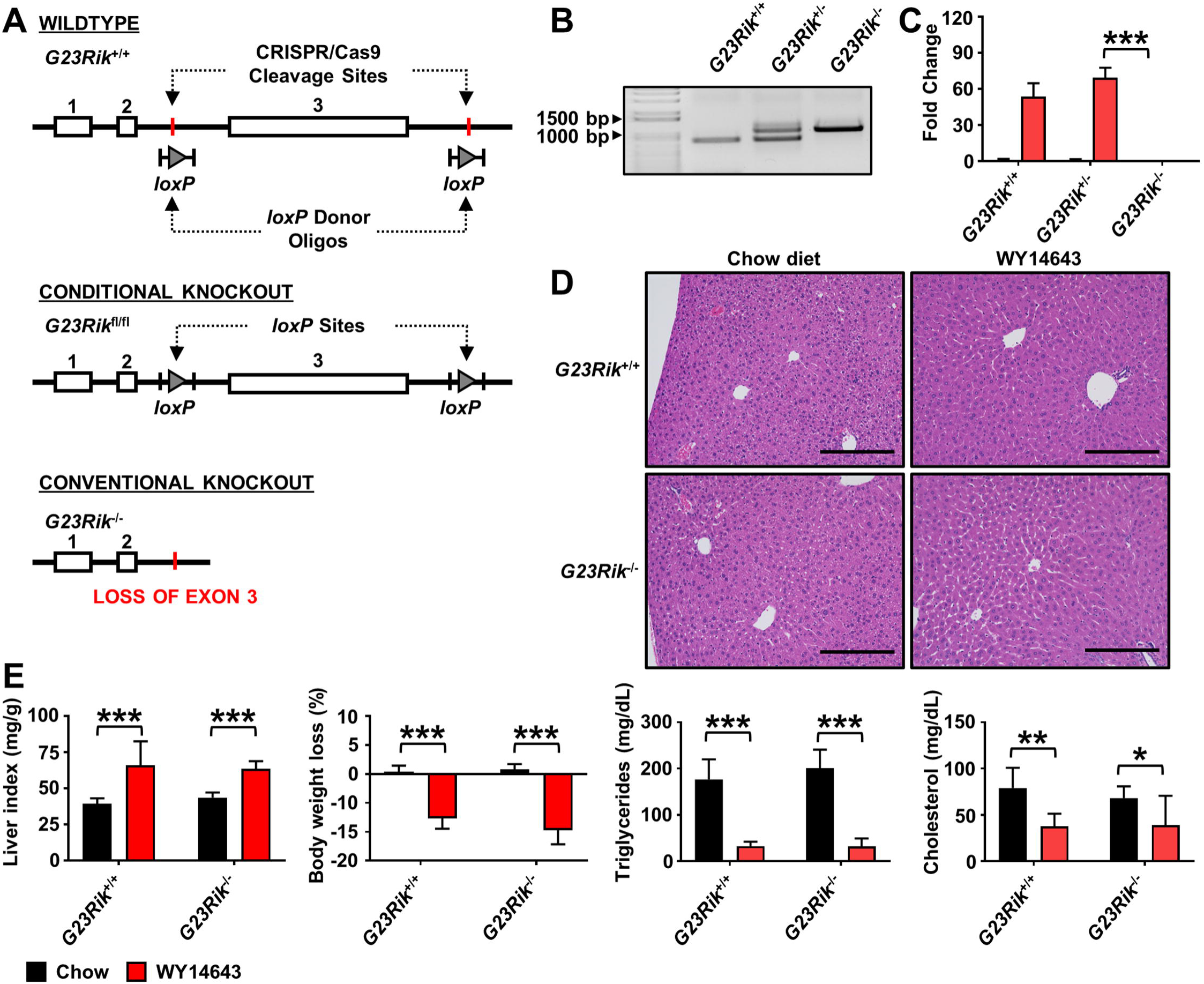
Generation of the *G23Rik*-null mouse line. Targeting strategy for generating a strand-specific *G23Rik*-null mouse line. **(A)** CRISPR/Cas9-mediated insertion of *LoxP* donor oligo between exon 3. Cre-mediated removal of *G23Rik* exon 3 blocks *G23Rik* expression. **(B)** *G23Rik* knockout mouse genotyping. **(C)** Analysis of *G23Rik* by qRT-PCR from livers of *G23Rik*^+/+^, *G23Rik*+/-, and *G23Rik*^-/-^ mice. **(D)** H&E staining of liver from *G23Rik*^+/+^ and *G23Rik*^-/-^ mice treated with chow diet or WY14643. **(E)** Liver index, body weight loss, triglycerides and cholesterol levels in liver of *G23Rik*^+/+^ and *G23Rik*^-/-^ mice. Each data point represents the mean ± SD for n = 5 liver samples. *P < 0.05; **P < 0.01; ***P < 0.001. Scale bars represents 20 nm (200x).

### Impact of G23Rik deficiency on the liver transcriptome

RNA-seq analysis was carried out to identify genes whose basal expression or response to WY14643 treatment was significantly different between *G23Rik*^+/+^ and *G23Rik*^-/-^ mouse liver. *G23Rik* was strongly upregulated by WY14643 in livers of wild-type mice but was not detected in *G23Rik*^-/-^ mice. In contrast, *Cyp4a14* mRNA, a classical WY14643-responsive gene, was significantly induced by WY14643 in livers of *G23Rik*^+/+^ and *G23Rik*^-/-^ mice (**Fig. 4A**). Further, a total of 16 genes whose basal expression was significantly altered in *G23Rik*^-/-^ mice were identified, 14 of which showed increased basal expression (**Fig. 4B**, *left*; **Table S2**). The most common pattern, seen for 9 of the 14 genes showing increased expression in the absence of *G23Rik*, was that WY14643 significantly reversed the increase in expression seen in *G23Rik*^-/-^ mice, but had little or no significant effect on gene expression levels in *G23Rik*^+/+^ mice (**Fig. 4B**, *middle vs. right*). This response pattern was particularly striking for four genes, the acute phase proteins serum amyloid A2 (*Saa2*), lipocalin 2 (*Lcn2*), orosomucoid 2 (*Orm2*), and serum amyloid A1 (*Saa1*), which were all strongly increased in livers of *G23Rik*^-/-^ mice as compared to *G23Rik*^+/+^ mice. *Lcn2* is up-regulated in response to inflammation, injury, and metabolic stress and protects from diet-induced fatty liver disease ^23, 24^. Saa1 potentiates fatty liver inflammation, while *Saa2* contributes to high density lipoproteins and cholesterol transport and *Orm2* functions as a tumor suppressor in liver cancer 23-28. The increased expression of these genes in livers of *G23Rik*^-/-^ mice was verified by qPCR (**Fig. 4C**). Interestingly, cytochrome P450 family 2 subfamily a member 4 (*Cyp2a4*) expression was induced by WY14643 to a much lower level in *G23Rik*^-/-^ liver compared to *G23Rik*^+/+^ liver, which is consistent with RNA-seq data and indicates a dependence on *G23Rik* for its full expression (**Fig. 4D**). In contrast, expression of seven other genes inducible by WY14643 was unchanged between *G23Rik*^+/+^ and *G23Rik*^-/-^ mice liver (**Fig. 4D**). These genes are indicative of the overall pattern, whereby loss of *G23Rik* did not have a significant effect on many genes responding to WY14643 treatment, with 617 genes induced by WY14643 and 521 genes repressed by WY14643 in both *G23Rik*^+/+^ and *G23Rik*^-/-^ mouse liver, and with no significant difference between models (**Table S2**).

**Fig. 4.**
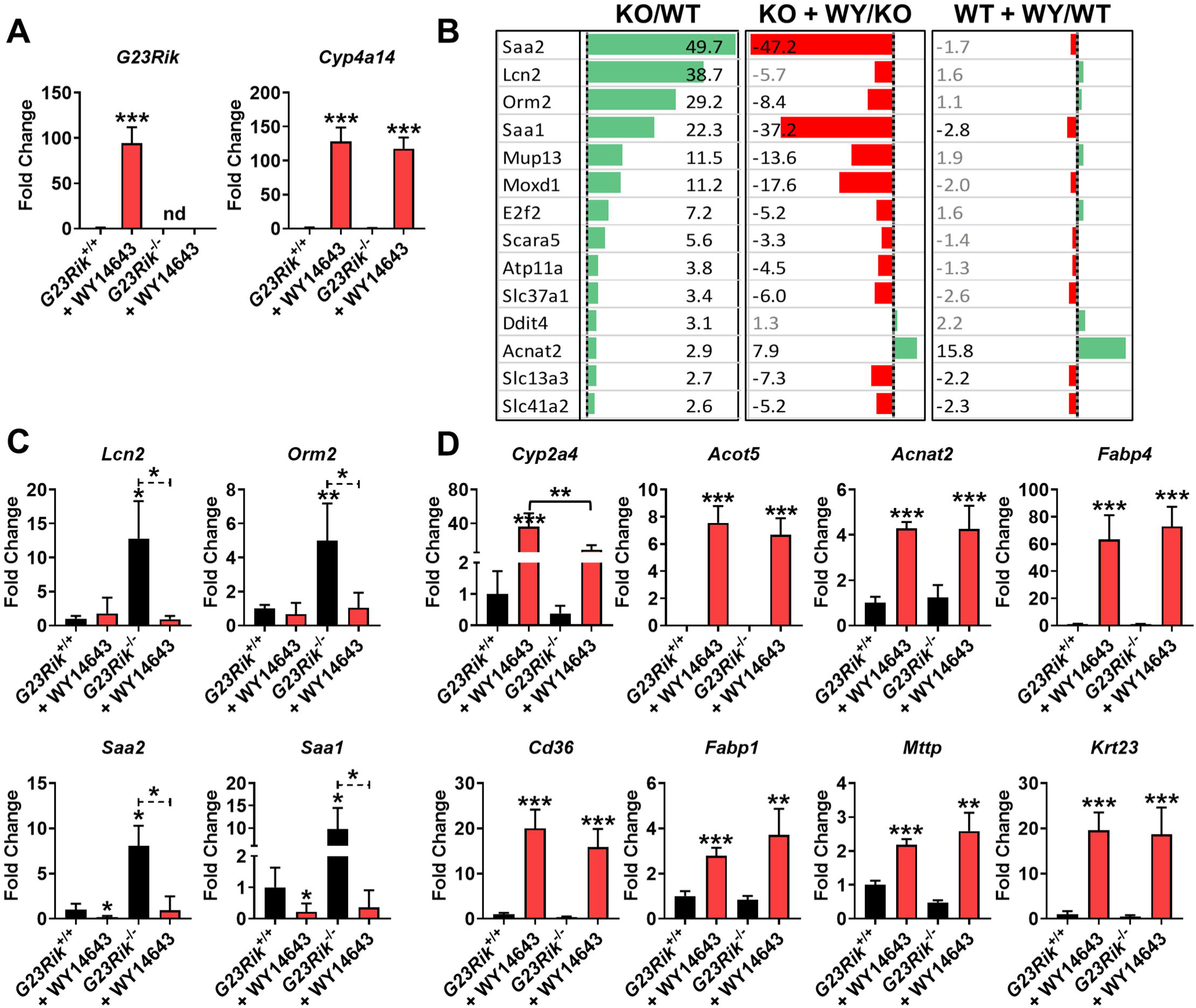
RNA-seq analysis from *G23Rik*^+/+^ and *G23Rik*^-/-^ mice liver. **(A)** Analysis of *G23Rik* and *Cyp4a14* mRNA by qRT-PCR from livers of *G23Rik*^+/+^ and *G23Rik-/-* mice treated with WY14643 for 48 h. **(B)** Liver RNA-seq analysis of 14 genes showing significant increases in expression in *G23Rik*^-/-^ compares to *G23Rik*^+/+^ mouse liver (*left* panel). 11 of the 14 genes showed a significant decrease in expression in livers of *G23Rik*^-/-^ mice treated with WY14643 for 48 h (*middle* panel). Very few genes showed a significant change in expression in livers of *G23Rik*^+/+^ mice given the same WY14643 treatment (*right* panel). Green, increase in expression; Red, decrease in expression, with the fold change values shown in black being significant at FDR< 0.05. See quantitative data in **Table S2. (C)** Analysis of *Lcn2, Orm2, Saa2*, and *Saa1* mRNA qRT-PCR from livers of *G23Rik*^+/+^ and *G23Rik*^-/-^ mice treated with WY14643 for 48 hrs. **(D)** Analysis of *Cyp2a4, Acot5, Acnat2, Fabp4, Cd36, Fabp1, Mttp*, and *Krt23* mRNA from qRT-PCR from livers of *G23Rik*^+/+^ and *G23Rik*^-/-^ mice treated with WY14643 for 48 h. Each data point represents the mean ± SD for n = 5 liver samples. *P < 0.05; **P < 0.01; ***P < 0.001.

### Loss of G23Rik potentiates lipid accumulation during physiological response to acute fasting

PPARα is activated by fasting, which facilitates metabolic remodeling that promotes the use of lipids as an alternate energy source and attenuates hepatosteatosis during fasting 1. To determine whether *G23Rik* protects against physiological fasting-induced liver damage, *G23Rik*^+/+^ and *G23Rik*^-/-^ mice were fasted for 24 h. Liver index and body weight loss were not different between fasted *G23Rik*^+/+^ and fasted *G23Rik*^-/-^ (**Fig. 5A**). Hepatic expression of *G23Rik* was significantly increased by fasting in *G23Rik*^+/+^ mice (**Fig. 5B**), although to a lower level than was seen following WY14643 treatment (c.f., **Fig. 4A**). Strikingly, liver lipid droplet accumulation was more widespread in fasting *G23Rik*^-/-^ mice than in fasting *G23Rik*^+/+^ mice (**Fig. 5C**). Oil Red O staining validated the greater lipid accumulation response to fasting seen in *G23Rik*^-/-^ liver (**Fig. 5C, Suppl. Fig. S2**). Serum total cholesterol (CHOL) levels and serum liver triglyceride (TG) levels were unchanged by fasting in either *G23Rik*^+/+^ or *G23Rik*^-/-^ mice. Serum non-esterified fatty acids (NEFA) were induced by fasting, but with no difference between *G23Rik*^+/+^ and *G23Rik*^-/-^ mice. In contrast, hepatic levels of CHOL, NEFA, and TG were significantly elevated in fasting *G23Rik*^-/-^ mice compared to *G23Rik*^+/+^ mice (**Fig. 5D**). Lipidomic profiling analysis showed that diacylglyceride (DG), TG, and phosphocholine (PC) accumulated in fasting *G23Rik*^-/-^ liver compared to fasting *G23Rik*^+/+^ liver (**Suppl. Fig. S3, Table S3**). Furthermore, hepatic expression of *Cd36*, a long chain fatty acid transporter and PPARA target gene ^29, 30^, was significantly elevated in fasting *G23Rik*^-/-^ mice compared to fasting *G23Rik*^+/+^ mice (**Fig. 5E**). Three other lipid metabolism-related genes (i.e., *Fabp1, Mttp*, and *Fabp4*) 11 which increased in response to WY14643 (**Fig. 4D**) did not show a fasting response in either *G23Rik*^+/+^ or *G23Rik*^-/-^ livers (**Fig. 5E**). Interestingly, the acute phase proteins *Lcn2, Orm2, Saa2*, and *Saa1* were all significantly elevated in livers of fed *G23Rik*^-/-^ mice when compared to fed wildtype animals (**Fig. 4A**). This suggests that in a fed state *G23Rik* expression may play a role in suppressing acute phase response in the liver. Expression of these four genes returned to wild-type levels after either WY14643 treatment or fasting in *G23Rik*^-/-^ mice (**Fig. 4D, Suppl. Fig. S4**). These genes are also known to be elevated in diet-induced fatty liver disease and other liver pathologies ^23, 24^. The role G23Rik plays during acute phase response in the liver is unknown and represents an interesting avenue for future research. Together, these results indicate that *G23Rik* may have regulatory role in the fasting-induced liver stress response.

**Fig. 5.**
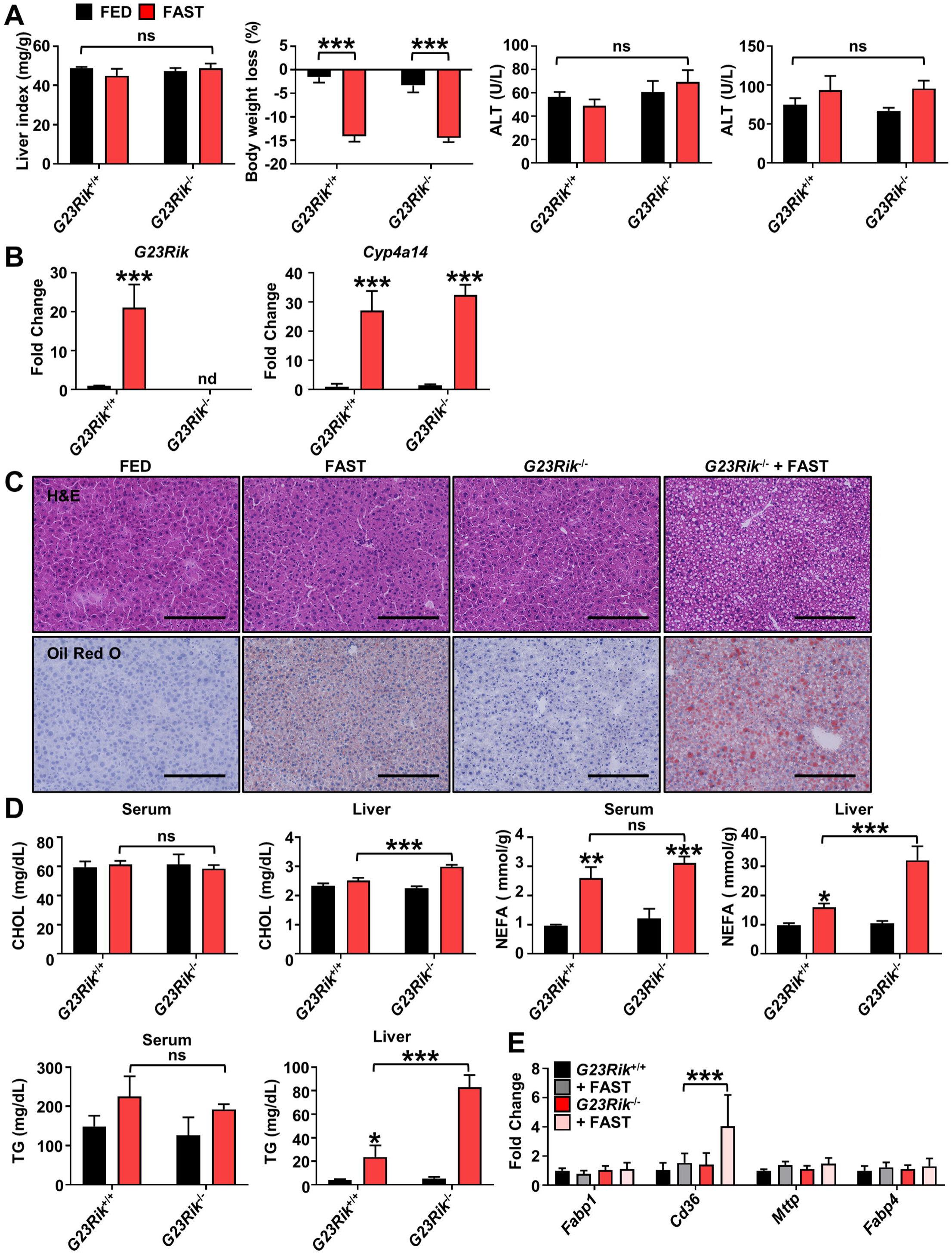
Loss of *G23Rik* potentiates lipid accumulation during physiological response to acute fasting. **(A)** Physiological endpoints from *G23Rik*^+/+^ and *G23Rik*^-/-^ mice after 24 hrs fasting. Liver indexes (mg liver/g body mass) and body weight loss in response to fasting. **(B)** *G23Rik* and *Cyp4a14* expression after 24 hrs fasting. **(C)** H&E and ORO staining of liver tissues after 24 hrs fast. Scale bars represents 100 nm (200x). **(D)** Cholesterol (CHOL), non-esterified fatty acid (NEFA) from serum, and triglycerides (TG) and levels from serum and liver tissues after 24 hrs fast. **(E)** Analysis of lipid metabolism-related gene mRNAs in livers of *G23Rik*^+/+^ and *G23Rik*^-/-^ mice after 24 hrs fast. Each data point represents the mean ± SD for n = 5 liver samples. *P < 0.05; **P < 0.01; ***P < 0.001. Scale bars represents 20 nm (200x).

## Discussion

The physiological alterations that accompany fasting impart metabolic flexibility that promotes several health benefits, including anti-inflammatory effects and prevention against metabolic diseases 31. Physiological PPARα activation by fasting is a key regulatory mechanism of lipid and glucose metabolism. Activation of PPARα modulates metabolic remodeling, inflammation, and cellular signaling pathways 32. Accumulating evidence indicates that PPARα activation suppresses inflammation in several disease models and tissues 33. An earlier study showed that the liver-enriched PPARα target lncRNA *Gm15441* attenuated liver inflammation and metabolic stress by suppression of the *Txnip*-mediated NLRP3 inflammasome pathway 11. However, the mechanism by which PPARα modulates metabolic stress is still unclear. The direct regulation of protein coding genes by PPARα is well studied, but there are many PPARα target lncRNAs whose functional or regulatory roles in liver are still unknown 11. PPARα regulated lncRNAs may contribute to physiological changes by promoting metabolic remodeling and gene expression through a variety of mechanisms.

RNA-seq analysis from liver treated with the specific PPARα agonist WY14643 identified more than 400 lncRNAs whose transcripts are significantly up regulated or down regulated in mouse liver 48 h after treatment with the potent PPARα agonist WY14643 11. One such lncRNA, *G23Rik*, was highly induced in WY14643-treated *Ppara*^+/+^ but not *Ppara*^-/-^ mouse liver. Basal expression of *G23Rik* was highest in muscle and BAT and relatively low in liver. However, following treatment with WY14643, *G23Rik* mRNA was significantly increased in liver but not in muscle or BAT, despite the expression of PPARα in those tissues 11. Following a single oral gavage of WY14643, hepatic *G23Rik* mRNA was rapidly induced within 1.5 h, with maximum expression seen after 6 h. The underlying PPARα regulatory mechanism controlling activation of the *G23Rik* was investigated. Motif analysis identified five potential PPARα binding sites within 5 kb of the *G23Rik* transcriptional start site, one of which was found to bind PPARα by ChIP and also to mediate transcriptional regulatory activity, as confirmed by luciferase reporter gene assays. Thus, lncRNA *G23Rik* is a direct PPARα target gene in the liver.

The activation of PPARα during fasting modulates fasting-induced metabolic stress responses, notably lipid accumulation and hepatosteatosis in mice 1. Lipid droplets appeared in livers of fasted *G23Rik*^+/+^ mice but were much more widespread in livers of fasting *G23Rik*^-/-^ mice. Moreover, liver CHOL, NEFA, and TG levels were significantly elevated by fasting in *G23Rik*^-/-^ mouse livers when compared to fasted wildtype animals. Expression of *Cd36*, a PPARA-regulated long chain fatty acid transporter and an important marker for metabolic-associated fatty liver disease ^29, 30^, was significantly elevated in livers of fasting *G23Rik*^-/-^ mice compared to *G23Rik*^+/+^ mice, which may contribute to the observed hepatosteatosis in *G23Rik*^-/-^ livers. Thus, *G23Rik* has a negative regulatory effect on *Cd36* expression under fasting conditions.

Expression of several liver damage marker genes, notably, *Lcn2, Orm2, Saa2*, and *Saa1*, was elevated in liver of *G23Rik*^-/-^ mice compared to *G23Rik*^+/+^ mice. Moreover, their expression was strongly suppressed by WY14643 treatment. *Lcn2* is an inflammatory protein elevated in inflammatory disease including fasting liver ^34, 35^. *Orm2*, an acute phage protein, is mainly produced in the liver where it is closely associated with cancer promoting pathways 27. *Saa2* and *Saa1* are acute immune response genes that promote inflammation-associated tumorigenesis in the colon 36. Expression of these liver damage marker genes is elevated in liver of fasting *Gm15441*-/- mice 11. The response of these four genes to *G23Rik* loss suggests that *G23Rik* has a modulatory effect on inflammatory-associated liver diseases through the inhibition of *Lcn2, Orm2, Saa2*, and *Saa1* expression. Further study is needed to clarify the underlying mechanism.

Interestingly, several prior gene expression studies that identified differential regulation of *G23Rik* also identified changes in *Lcn2* expression suggesting these two genes may be functionally related. *G23Rik* was identified as one of the top 50 differentially regulated genes in the pancreatic islet cells of mice lacking the short chain fatty acid receptor *Ffar3* 15. That study found that *Ffar3* signaling regulates glucose stimulated insulin secretion. *Lcn2* was also identified as one of the top 50 most differentially expressed genes 15. Another study found *G23Rik* was upregulated over seven-fold and was one of the top 20 differentially expressed lncRNAs in the brains of mice infected with the rabies virus (RABV) 37. Again, *Lcn2* was one of the top differentially expressed mRNA identified in that study. The relationship between *G23Rik* and *Lcn2* represents an interesting focus of future research.

In summary, the PPARα-regulated hepatic lncRNA *G23Rik* is a potential metabolic regulator that is specifically up regulated in liver in response to PPARα activation. Fasting induced lipid accumulation in *G23Rik*^-/-^ mouse liver when compared to fasted wild type mice. Liver TG, CHOL, and NEFA levels were significantly elevated and associated with induction of *Cd36*, an important lipid metabolism-related gene. Together, these results indicate that hepatic lncRNA *G23Rik* can prevent physiological fasting-induced inflammation and lipid accumulation and represents a novel therapeutic target for the treatment of inflammatory disorders.

## Supporting information

Suppl Figures listing

## Abbreviations

Acot1: encoding acyl-CoA thioesterase 1
Acox1: encoding acyl-Co-A oxidase 1
ALT: alanine aminotransferase
AST: aspartate aminotransferase
BAT: brown adipose tissue
Blnc1: brown fat lncRNA 1
CHOL: total cholesterol
Cyp2a4: cytochrome P450 family 2 subfamily a member 4
DG: diacylglyceride Ffar3, free fatty acid receptor-3
G23Rik^-/-^: G23Rik knockout mice
G23Rik: lncRNA 3930402G23Rik
HIF1α: hypoxia-inducible factor 1α
Lcn2: lipocalin 2 Long non-coding RNAs
(lncRNAs): 
NEFA: non-esterified fatty acids
Orm2: orosomucoid 2
Ppara^-/-^: PPARα knockout mice PPARα; peroxisome proliferator-activated receptor α
PPRE: peroxisome proliferator response elements
Saa1: serum amyloid A1
Saa2: serum amyloid A2
TG: triglycerides
UPLC-ESI-QTOFMS: UPLC-electrospray ionization-quadrupole time-of-flight mass spectrometry.

## Acknowledgements

We thank Linda G. Byrd and John R. Buckley for technical assistance with the mouse studies. This work was funded by the National Cancer Institute, National Institutes of Health Intramural Research Program and by NIH grant ES024421 (to DJW).

