## Supplementary material for "Long non-coding RNA G23Rik attenuates fasting-induced lipid accumulation in the liver": Suppl Figures listing

**Fig. S1. PPARα ChIP-seq peak of upstream of *G23Rik*.**

PPARα ChIP-seq read peaks from agonist (GW7647)-treated mouse liver.

**Fig. S2. Oil Red O staining in liver of fed or fasting *G23Rik*^+/+^ and *G23Rik*^-/-^ mice.**

Representative ORO staining of liver tissues after a 24 h fed or fast in *G23Rik*^+/+^ and *G23rik*^-/-^ mice (n = 5, 3 images/mouse). Scale bars represents 100 nm (×200)

**Fig. S3. Lipidomics profiling analysis in *G23Rik*^+/+^ and *G23Rik*^-/-^ mice under fed or fasting conditions.**

(A) PCA score plots for ESI+ mode. (B) Relative intensity of lipid markers in liver of *G23Rik*^-/-^ mice and WT mice. * FDR adjusted p-value <0.05

**Fig. S4. Lipid metabolism-related gene expression in *G23Rik*^+/+^ and G23Rik^-/-^ mice under fed or fasting conditions.**

Hepatic expression of lipid metabolism-related genes (*Orm2*, *Saa1*, *Saa2*, and *Lcn2*) in *G23Rik*^+/+^ and *G23Rik*^-/-^ mice fed or fasted for 24 hours as determined by qRT-PCR. Each data point represents the mean ± SD for n = 5 liver samples. Each data point represents the mean ± SD for n = 5 liver samples. *P < 0.05; **P < 0.01.

**Table S1**. Sequences in this study.

| **Name** | **Sequence (5’ → 3’)** |
| --- | --- |
| Acnat2 F | GATTCTGAATGGCAGGGAGGAA |
| Acnat2 R | GGCTCATCCACAAGGGCATT |
| Cd36 F | GATTAATGGCACAGACGCAGC |
| Cd36 R | CAGATCCGAACACAGCGTAGA |
| Cyp2a4 F | TGGGCTGCATGAGGTTAAAGG |
| Cyp2a4 R | TCCTCTGCAAGTCTACGAAGT |
| Cyp4a14 F | CCTGACTTTCTTTCGCCTGC |
| Cyp4a14 R | TGATCACTCCATCTGTGTGCT |
| Ddit4 F | GGTCTGCAGCCAGAGAAGAG |
| Ddit4 R | TCCAGGTATGAGGAGTCTTCC |
| E2f2 F | ACGGCGCAACCTACAAAGAG |
| E2f2 R | GTCTGCGTGTAAAGCGAAGT |
| Fabp1 F | AGTCAAGGCAGTCGTCAAGC |
| Fabp1 R | ATGTCGCCCAATGTCATGGT |
| Fabp4 F | CATAACCCTAGATGGCGGGG |
| Fabp4 R | CGCCTTTCATAACACATTCCACC |
| GAPDH F | GACTTCAACAGCAACTCCCAC |
| GAPDH R | TCCACCACCCTGTTGCTGTA |
| G23Rik F | ATGGACGTGAAGAACCAGGAA |
| G23Rik R | CAGATCTGGGGGTGATGTGC |
| Lcn2 F | TCTGTCCCCACCGACCAAT |
| Lcn2 R | GGAAAGATGGAGTGGCAGACA |
| Mttp F | CCCACTCAGGCAATTCGAGA |
| Mttp R | TGATGGAGGGGGAGTTCACA |
| Mup13 F | GATGCTGTTGCTGCTGTGTT |
| Mup13 R | TGCCATTCCCCATTAATCTTTTCT |
| Orm2 F | ATTGGTGCGGCTGTCCTAAA |
| Orm2 R | ACACAGTGGTCATCTATGGTGT |
| Saa1 F | GACACCAGGATGAAGCTACTCACC |
| Saa1 R | CCCCTTGGAAAGCCTCGTGA |
| Saa2 F | ACACCAGCAGGATGAAGCTACT |
| Saa2 R | CCTTGGAAAGCCTCCCCAATA |
| Slc37a1 F | CTCTGTGGCCAACGCAGT |
| Slc37a1 R | GGCTGATCTCTGCGGGC |
| ChIP Acot1 F | CACCGGAGTCACCTGATAGAGTC |
| ChIP Acot1 R | GCCAGGGTGCACAGACTTT |
| ChIP Acox1 F | CGGAAACCAGAAGGGAATG |
| ChIP Acox1 R | TAGCCAACGACAATGAACC |
| ChIP Block A F | GGCAACGATCCCTCCCCTAT |
| ChIP Block A R | AGCCCCGGCTCAATGAAATG |
| ChIP Block B F | CACCAAATTCGTAACGTAGTGT |
| ChIP Block B R | CAATGGCCTTACATGTTTTAACTCT |
| ChIP Block C F | CCGTGGAAATTCATTGACGCA |
| ChIP Block C R | GGAAGGCATGTCAACACACG |
| ChIP Block D F | GGCACATATTGCCGTTGCTT |
| ChIP Block D R | AGTGGGATCTGTGAGTCCCC |
| ChIP Block E F | AGGCTTCACATGAGTGTGCT |
| ChIP Block E R | GCCTCCAGACGTAAACGCAT |
| G23Rik 5' loxP F | CTACTCCAGCCAACACCAGG |
| G23Rik 3' loxP R | GCCTGCACAAGGGGGAGGTC |
| G23Rik 3' loxP R | TCCCCAGCTCACTGGAAGAAGC |
| G23Rik  Geneblock A | ATTCTCTGGCCTAACTGGCCGGTACCAGGTTGGGAAATCAGCTTGTCCACAGAGCCAGATCCCCCTGGCAACGATCCCTCCCCTATAGACAGAGAAAGGCAAAGCCCCTGAGAGGAGCTCAGCGGAAAGTACATTTCATTGAGCCGGGGCTTGGTTACAAAATAGATAAACTTGCACCGTTTCCAGGCCAACGGGATTCAAGCTGCCGAGATGTTGACCAAGTTGGAAGCTTGGCAATCCGGTACTGTTGG |
| G23Rik  Geneblock B | ATTCTCTGGCCTAACTGGCCGGTACCGACTTTGGTCAATCTCTTTGATGGATCCTTCTAACTTTACTCAGTGTCTTGATTGTGACTAATTGATGCCTTAAGACACCAAATTCGTAACGTAGTGTACTGTTCACCAGCACAGCACTGCAGTTATAAAGCATAACATGTTTGTTTACAAAACAAATAGAGTTAAAACATGTAAGGCCATTGCATACCTGGATTCATATAAGCTTGGCAATCCGGTACTGTTGG |
| G23Rik  Geneblock C | ATTCTCTGGCCTAACTGGCCGGTACCCCAAGTATCGTGTGTTGACATGCCTTCCTGGGCCTCCCCAAGGCCCTGTCCTCAAGAACAATGTAACCTGGATTAGGAAGACAGGTCAGATGATTCTAAATGATGAAACTGCATAGGTCAGAGCAGCAAGAGGATAAGGCCAGGTAAGGTGAGGCCAGGTAAGGGGGGTGTTTCCTGCTGTCCTGGGACGTGGAAACTTAAAGCTTGGCAATCCGGTACTGTTGG |
| G23Rik  Geneblock D | ATTCTCTGGCCTAACTGGCCGGTACCGCTTCATAGCAACATCCTTCCCACTCCAGATGTGTAAAAATAATAACCTGATAGCCCCGGGGACTCACAGATCCCACTCTGTTGTCTATGTCTCAGTGCAAGTTCATTTGTGGTTTTAGCATCTATGCAATAACATTAACAAAATCCAGAAGCCATATGCTGCTTTCACCTCTGCCTAAAATGTCTGAAGTGGAGGAGGCAAGCTTGGCAATCCGGTACTGTTGG |
| G23Rik  Geneblock E | ATTCTCTGGCCTAACTGGCCGGTACCTGAGATAAAATAGTATGTCTCAGTTGTTGCATGTCACCTTTTAAAGCACAAAGTTCTTAATGCGTTTACGTCTGGAGGCCAATTAGAGAAAACAGTGGCTCCATGACCCAGTGAGTGGCTGGGCCCTTCTACATTGTTCCATTCTGAGTTCTGGGAGGCTTCCAGTCCTCAGAGTCAGGATGAGGAGCTTGGAGGGAAGCCATAGGTCAACATATGCCCATCACTGTGGTTATAAATAGAAGTTAGCTGGAAGCTTGGCAATCCGGTACTGTTGG |
| G23Rik Upstream Oligo donor | gagctgcagtcctgtcacccaaggaactgaactgagtagccaatggctccatgccactgacaatttatggctcgagataacttcgtatagcatacattatacgaagttatcggaggcaaggattctacttggacatagcaggagcaatactaagccagtgccacctcccagtgaggcctc |
| G23Rik Downstream Oligo donor | gcaaataagatctgtgtagaggcttgcaagtccccactgtggcccaggcggcagggtttctcaggagcgctcgagataacttcgtatagcatacattatacgaagttattatggggtgtgatcaggctttgcatcaggaatgggaaagaaccacttcaccccacaaaatgtgtagttag |

*Red text denotes sequence that is PPARa binding elements

**Table S2**. Differentially expressed genes identified by RNAseq analysis. See separate Excel file.

**Table S3**. Identified lipids that were significantly accumulated in fasting *G23Rik*^-/-^ liver.

| ID | m/z | RT (min) | possible adduct |
| --- | --- | --- | --- |
| TG 58:10 | 909.7351 | 17.41 | [M+H-H2O]+ |
| TG 56:9 | 901.7304 | 17.03 | [M+H]+ |
| TG 56:3 | 930.8512 | 17.85 | [M+NH4]+ |
| TG 54:5 | 898.8927 | 17.16 | [M+NH4]+ |
| TG 54:2 | 904.8358 | 17.84 | [M+NH4]+ |
| TG 52:3 | 874.7891 | 17.26 | [M+NH4]+ |
| Fragment of TG 52:3 | 575.5054 | 17.23 |  |
| TG 52:1 | 878.82 | 17.84 | [M+NH4]+ |
| TG 50:1 | 850.7891 | 17.51 | [M+NH4]+ |
| DG 36:2 | 603.5364 | 17.54 | [M+H-H2O]+ |
| DG 36:1 | 605.5518 | 17.84 | [M+H-H2O]+ |
| DG 34:2 | 575.5049 | 16.94 | [M+H-H2O]+ |
| PC 35:4 | 785.5729 | 14.29 | [M+NH4]+ |
