## Supplementary figures and images for "Long non-coding RNA G23Rik attenuates fasting-induced lipid accumulation in the liver"

### Supplemental Figures

#### Slide 1
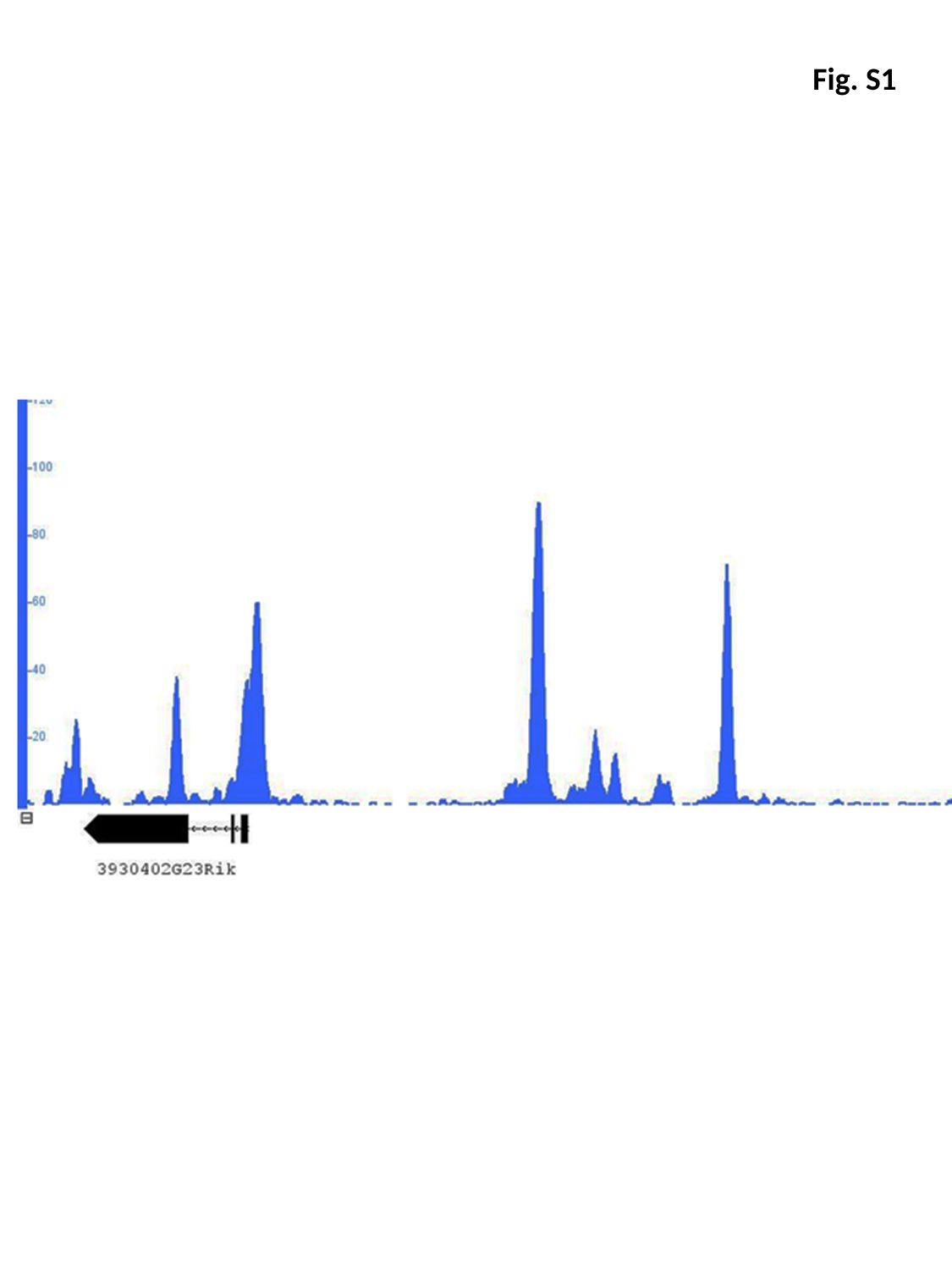

Fig. S1

#### Slide 2
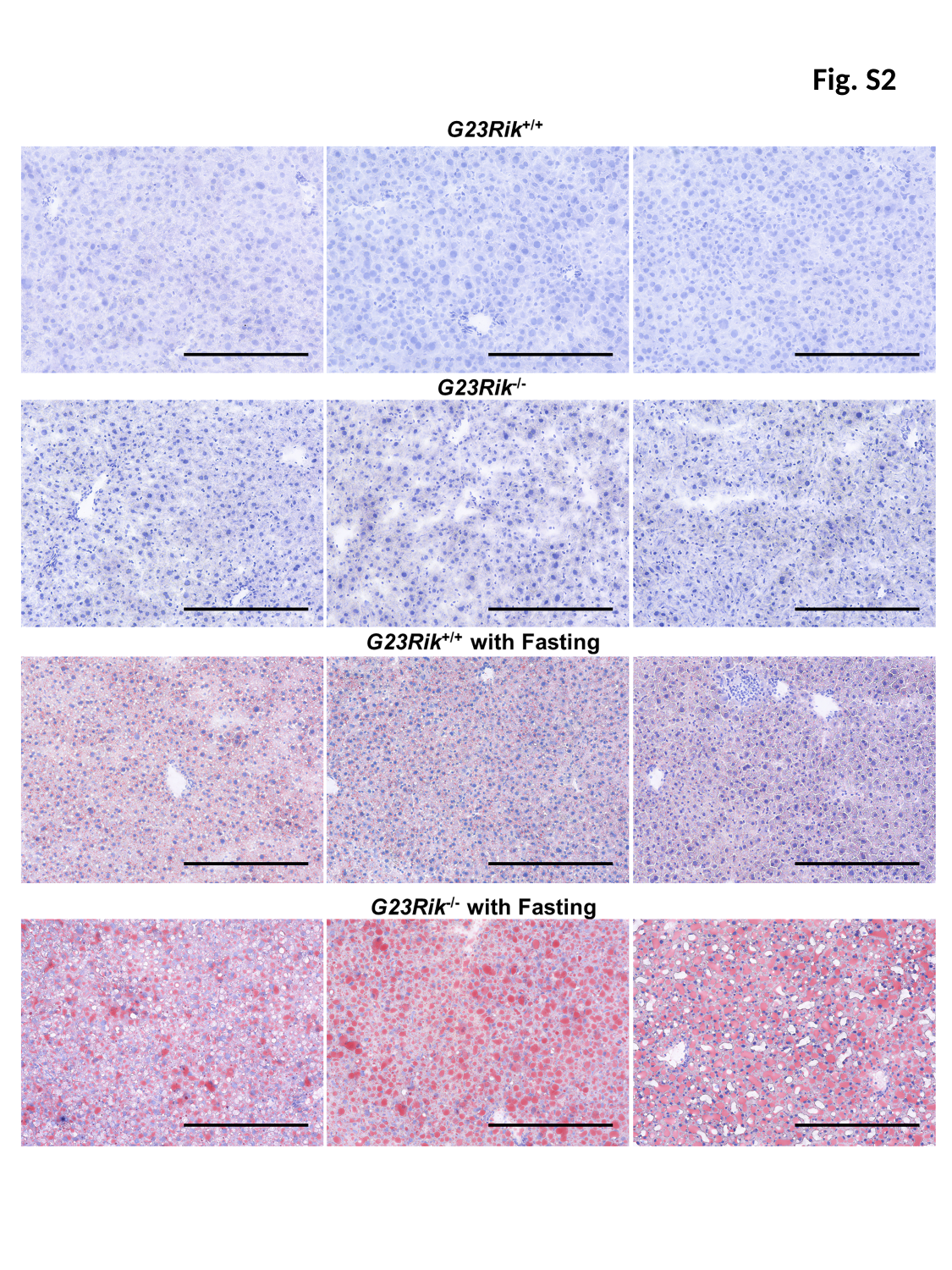

Fig. S2

#### Slide 3
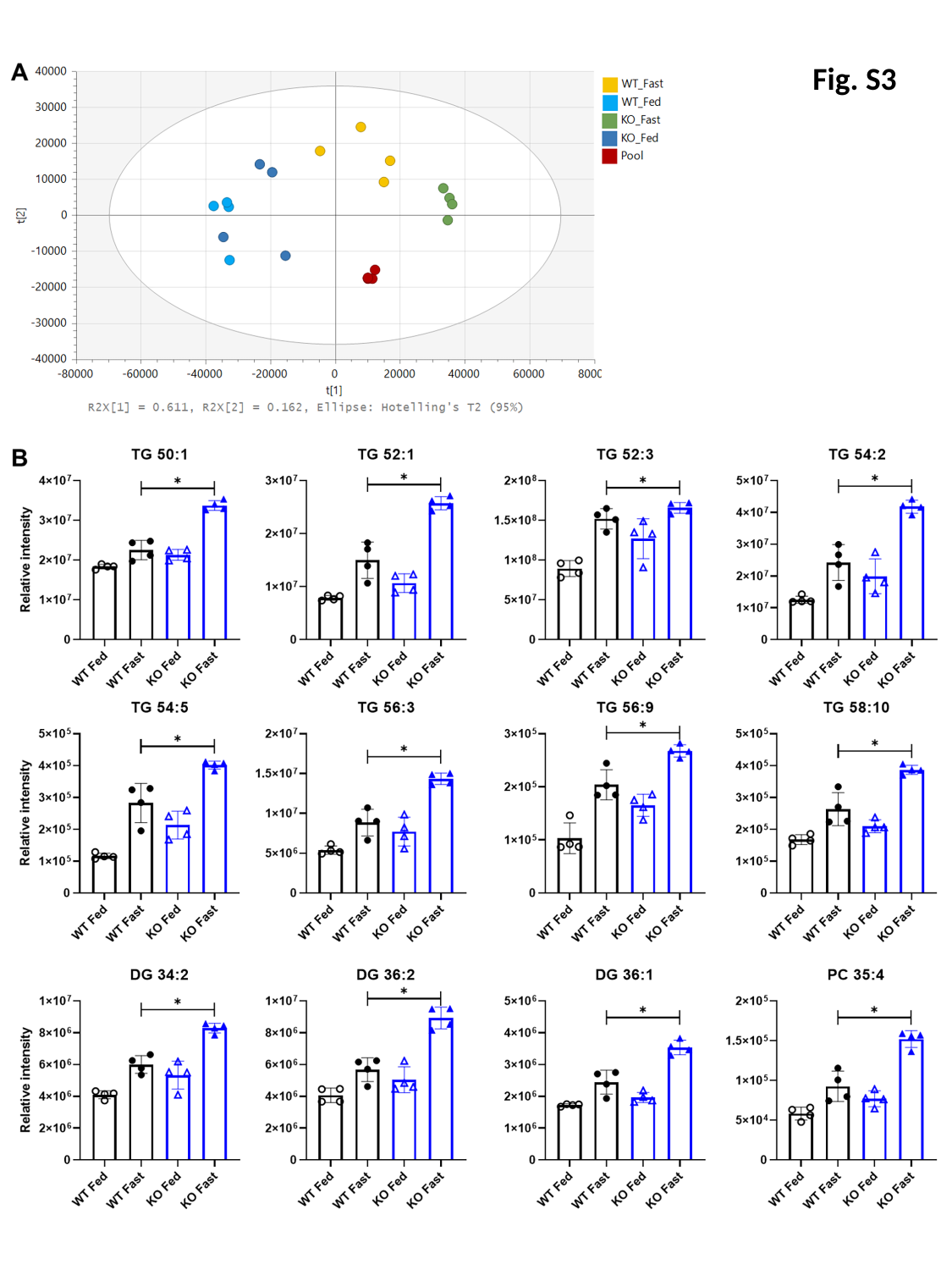

Fig. S3

#### Slide 4
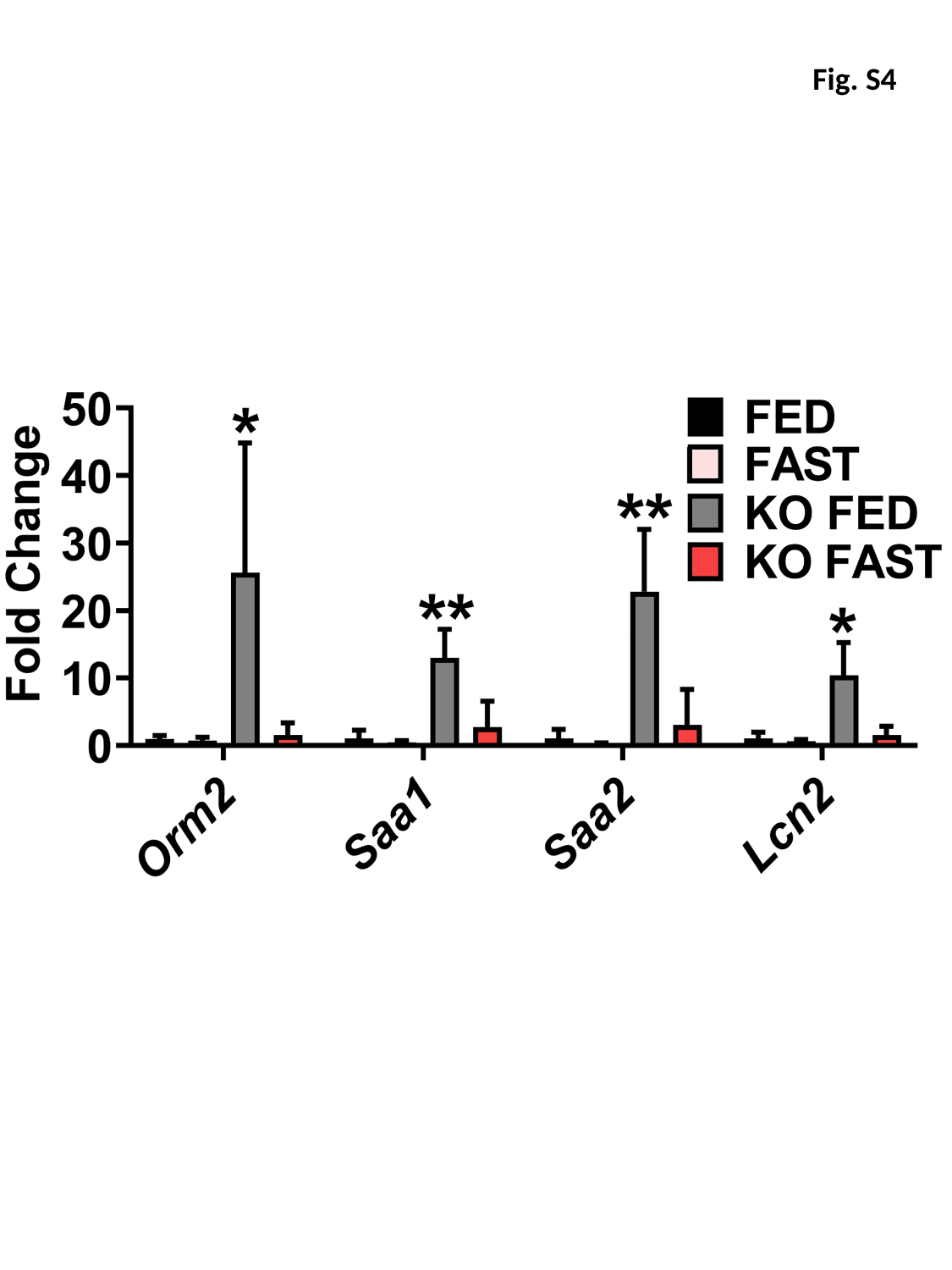

Fig. S4
